# The transcriptional repressor LjTIE1 constrains cytokinin signaling to balance infection and organogenesis during root nodule symbiosis

**DOI:** 10.64898/2025.12.19.694972

**Authors:** Maxim Yorick Limbächer, Manuel Frank

**Affiliations:** Department of Molecular Biology and Genetics, Aarhus University, Universitetsbyen 81, DK-8000 Aarhus C, Denmark.; Institute of Plant Biology and Biotechnology (IBBP), University of Münster, Schlossplatz 8, D-48143 Münster, Germany.

## Abstract

- Cytokinin (CK) is a key regulator of root system architecture including primary root (PR) and lateral root (LR) development. During root nodule symbiosis in legumes, CK promotes nodule organogenesis in the root cortex while simultaneously suppressing infection thread (IT) formation in the epidermis, yet the molecular mechanisms enabling this spatial specificity remain incompletely understood. TCP INTERACTOR CONTAINING EAR MOTIF PROTEIN (TIE) proteins negatively regulate CK signaling in Arabidopsis roots, but whether legume orthologs modulate symbiotic CK responses remains unknown.
- We characterized Lotus *tie1* loss-of-function mutants through root and nodulation assays, transcriptomics, confocal microscopy of CK signaling reporter TCSn, and ethylene quantification.
- Lj*TIE1* expression was induced by CK and Nod-factor, *tie1* mutants exhibited reduced PR length, LR density, IT formation, and nodule number alongside elevated ethylene emission - phenotypes fully rescued by ethylene biosynthesis inhibition. Despite enhanced CK signaling (confirmed via TCSn reporter and transcriptomics), *tie1* roots formed no spontaneous nodules but displayed hypernodulation under exogenous CK treatment.
- LjTIE1 provides temporal control of symbiosis by dampening CK signaling after nodule initiation, revealing a regulatory layer that prevents constitutive nodulation despite elevated CK perception.

## Introduction

As sessile organisms, plants dynamically adapt their shoot and root system to the environment maximising their chances for survival. The root system is crucial for the uptake of nutrients from the surrounding soil and anchors the plant in it. Phytohormones such as cytokinin (CK) - an adenine-derivative - regulate root system development and integrate external environmental cues. CKs are crucial for maintaining root apical meristem function, development of lateral roots, uptake of nutrients, and establishment of plant-microbe interactions (Sakakibara, 2006; Cortleven *et al.,* 2019; Lin *et al.,* 2020). N_6_-isopentenyladenine (iP) and *trans*-zeatin (*t*Z), synthesized via ISOPENTENYL TRANSFERASEs (IPTs) and Cytochrome P45O monooxygenases (CYP735As) (Takei *et al.,* 2004; Kiba *et al.,* 2013), and subsequently activated by LONELY GUYs (LOGs) (Kurakawa *et al.,* 2007), represent the primary bioactive forms regulating root development. Bioactive CKs may be degraded by CK OXIDASE/HYDROGENASEs (CKXs) (Werner *et al.,* 2001) or may be inactivated via N-glucosylation. In their bioactive form, CKs are perceived by HISTIDINE KINASEs (HKs) which are activated by autophosphorylation (Inoue *et al.,* 2001; Ueguchi *et al.,* 2001; Yamada *et al.,* 2001). Subsequently, HKs phosphorylate HISTIDINE PHOSPHOTRANSFER PROTEINs (HPTs/HPs) (Hutchison *et al.,* 2006). HPTs ultimately phosphorylate type B RESPONSE REGULATORS (RRBs) which accumulate in the nucleus and activate CK-responsive gene expression (Imamura *et al.,* 1999; Sakai *et al.,* 2000, 2001). Two well-studied mechanisms repressing CK signaling are facilitated by HPTs lacking the canonical phosphoryl moiety acceptor histidine residue and by type A response regulators (RRAs) (Brandstatter & Kieber, 1998). While CK’s role in root architecture is well-established, legume-rhizobia symbiosis reveals how a single hormonal signal can drive distinct developmental outcomes in adjacent tissue layers (Lin *et al.,* 2020). As a result of Nod-factor perception and activation of the symbiotic signaling pathway, bioactive CKs are synthesized in Lotus susceptible zones and synthesis genes such as *IPT2, IPT3, IPT4* and *LOG4* are upregulated (Reid *et al.,* 2017). Synthesized CKs bind to the LOTUS HK1 (LHK1), activate the CK signaling pathway which ultimately regulates nodule organ formation through dedifferentiation of cortex and pericycle cells (Murray *et al.,* 2007; Tirichine *et al.,* 2007). Therefore, supplementation with CKs such as the synthetic 6-Benzylaminopurine (BAP) is sufficient to induce spontaneous nodulation in the absence of rhizobia in Lotus (Heckmann *et al.,* 2011) and many other legume species (Gauthier-Coles *et al.,* 2018). Similarly, point mutations in the CHASE domain of LHK1 resulting in constitutive CK signaling induce spontaneous nodules (Tirichine *et al.,* 2007). Downstream of CK, NODULE INCEPTION (NIN), NODULATION SIGNALING PATHWAY 1 (NSP1) and NSP2 regulate nodulation (Kaló *et al.,* 2005; Smit *et al.,* 2005; Heckmann *et al.,* 2011). While CK is a positive regulator of nodulation, its application and Nod-factor induced CK synthesis inhibit root hair colonization through LHK1 (Murray *et al.,* 2007; Tirichine *et al.,* 2007). *Ihk1* roots therefore display a hyperinfection phenotype while being impaired in nodule organ formation (Murray *et al.,* 2007). This spatial separation indicates that CK’s opposing effects on infection versus organogenesis require cell-type-specific regulatory components beyond the central HK receptor. However, the transcriptional modulators that confer tissue-specific CK responses during symbiosis remain poorly characterized. For nodulation and root hair infection, CK action ultimately requires the synthesis of the gaseous phytohormone ethylene (Lin *et al.,* 2020). It is synthesized right after rhizobial perception and acts as a negative regulator for both processes (Reid *et al.,* 2018). The application of the ethylene biosynthesis inhibitors Aminoethoxyvinylglycine (AVG) or Ag+, and an impairment in ethylene perception enhances nodulation and infection in a variety of legumes such as pea (Goodlass & Smith, 1979; Lorteau *et al.,* 2001), Medicago (Peters & Crist-Estes, 1989; Penmetsa & Cook, 1997) and Lotus (Heckmann *et al.,* 2011; Reid *et al.,* 2018). ETHYLENE RESPONSE FACTOR-ASSOCIATED AMPHIPHILIC REPRESSION (EAR) MOTIF-CONTAINING PROTEIN TEOSINTE BRANCHED 1/CYCLOIDEA/PROLIFERATING CELL FACTOR (TCP) INTERACTOR CONTAINING EAR MOTIF PROTEIN (TIE) proteins are transcriptional repressors regulating shoot and root development. They contain a nuclear localization signal (NLS), a SPL-motif/helix region and an EAR motif (Tao *et al.,* 2013). In Arabidopsis shoots, TIEs interact and suppress TCP transcription factors by forming a complex with TOPLESS (TPL)/TOPLESS-RELATED (TPR) corepressors (Tao *et al.,* 2013). TIEs moreover interact with TIE1-ASSOCIATED RING-TYPE E3 LIGASE1 (TEAR1) leading to their ubiquitination and subsequent degradation (Zhang *et al.,* 2017). Recently, AtTIE1 and AtTIE2 were found to be crucial for CK signaling. Both genes are induced upon CK perception and the translated proteins impair RRB function by forming a TIE-RRB-TPL/TPR complex (He *et al.,* 2022). Consequently, *tie1tie2* loss-of-function double mutants have increased CK signaling with an increased emission of ethylene resulting in an impairment of primary and lateral root growth (He *et al.,* 2022). While the role of AtTIEs during CK-dependent root development is well described, it is unclear if and how homologues in legumes like *Lotus japonicus* impact root and nodule development. In this study, we identified one homologous *TIE* gene in Lotus, which we named Lj*TIE1.* Like in Arabidopsis, Lj*TIE1* was regulated by CK and a loss of Lj*TIE1* resulted in shorter roots, a reduced lateral root density and a pronounced ethylene synthesis as a result of increased CK signaling. Compared to the *LHK1* gain-of-function mutant *snf2,* the enhanced CK signaling in *tie1* mutants did not result in spontaneous nodulation or induction of symbiotic signaling in the absence of rhizobia. Our study therefore highlights the importance of signaling specificity: enhanced CK signaling alone is insufficient for nodule organogenesis without appropriate symbiotic perception, revealing a regulatory layer between CK perception and nodule development that differs from artificial CK treatments or constitutive receptor mutations.

## Materials and Methods

### Plant material and growth conditions

*Lotus japonicus* seeds were scarified with sand paper and sterilized in a 1% sodium hypochlorite solution for 15 minutes. After three washes with sterile water, seeds were placed on water-soaked filter paper in square plates and germinated. Three days after germination, seedlings were transferred to square plates with 1.4% Agar Noble slopes containing O.25x Broughton & Dilworth (B&D) medium covered with filter paper. A metal bar with 3-mm holes for roots was inserted at the top of the agar slope. For genetic studies in Lotus, LORE1 lines *tie1-1* (30090728), *tie1-2* (30139000) (Malolepszy *et al.,* 2016), and *snf2* (Tirichine *et al.,* 2007) were used. Treatments with AVG and BAP were started from the start of germination until quantification of root characteristics.

### Nodulation assays and infection thread quantification

For nodulation assays, seedlings were inoculated with 500 µL of either water (control) or *Mesorhizobium loti* strain R7A (R7A, OD600 = 0.02) along the length of the root. Pink and white nodules were quantified at the indicated time points after inoculation. For infection thread (IT) quantification, dsRed-labeled R7A was used. ITs were counted at the primary root. To determine IT density, primary root (PR) length was measured using ImageJ® (Abramoff *et al.,* 2004), and IT counts were normalized to PR length to account for growth variation.

### Ethylene measurements

Ethylene measurements were performed as described in (Reid *et al.,* 2018). In brief, two 3-day-old seedlings were transferred to 5 mL glass GC vials containing filter paper wet with 750 mL nitrate-free O.25x B&D nutrients. After two days of adjustment to vials, seedlings were inoculated with 250 µL of either water or *M. loti* R7A (OD600 = 0.02) along their roots. One day later, GC vials were capped with synthetic stoppers from Vacuette Z No-Additive tubes (Greiner Bio-One, product no. 455001). Ethylene accumulation over the following 24 h was measured using an ETD-300 laser-based detection system (SensorSense).

### Hairy root transformation

*pIV1O-TCSn::tYFPnls* containing Agrobacterium rhizogenes ARI 193 (Reid *et al.,* 2017) were introduced into wild-type, *tie1-1* and *tie1-2* seedlings by piercing the hypocotyl with a narrow needle. Three weeks later, the non-transgenic primary root of infected seedlings was cut, and plants exhibiting hairy roots were placed into pots containing lightweight expanded clay aggregate (LECA). After 9 days, cells walls of hairy roots were stained with a 0.5 mg/mL Calcofluor white solution for 2 minutes and hairy roots were imaged immediately after.

### Confocal microscopy and nuclei intensity quantification

Confocal microscopy was performed with a Zeiss LSM780 microscope. Following excitation/emission [nm] settings were used: (i) Calcofluore white-stained cell wall: 405/420-505, (ii) DsRed and mCherry: 561/580-660, (iii) tYFP:: 514/517-560. tYFP signal was detected with 1.8 % laser power and a gain of 650 for all hairy roots to ensure comparability between roots. Laser power and gain settings were optimized on wt control roots and held constant to allow inter-genotype comparison, even if signal saturation occurred in enhanced-signal *tie1* samples. Nuclei quantification was performed using ImageJ®. Maximum projections were converted to 16-bit pictures, 10 units of background subtracted using the Math function and the threshold adjusted until the majority of nuclei around the meristem were red without overlapping. Then, pictures were converted into binaries and the function “Watershed” applied to separate overlapping nuclei. Nuclei were identified by the software using the “Analyze Particles” function and the created ROI mask was used on the original maximum projection to measure nuclei intensities using the “Measure” function in the ROI Manager.

### RNA-sequencing and data analysis

For RNA-sequencing, whole roots of seedlings were harvested 3 dpi with either water or R7A. Total RNA was isolated using the InviTrap® Spin Plant RNA Mini Kit (Invitek Diagnostics) following the manufacturer’s instructions. Library preparation (poly-A enrichment) and sequencing (Novaseq X Plus, PEI50) was conducted by Novogene. RNA-seq data analysis was performed with R® 4.5.1 in RStudio®. Preprocessing of paired-end reads including trimming were performed with Fastp (Chen *et al.,* 2018) and quality control performed with FastQC. Reads mapped against the Lotus Gifu 1.3 reference genome using STAR (Dobin *et al.,* 2013) and aligned reads were counted with the STAR Genecount function. DEGs were extracted using DESeq2 (with BH-corrected p < 0.05) (Love *et al.,* 2014). Raw data are deposited at the Sequence Read Archive (SRA4) with the BioProject ID PRJNA1333668.

### Sequence alignment

TIE amino acid sequences were taken from Lotus Base (Mun *et al.,* 2016) and Aramemnon (Schwacke *et al.,* 2003). Sequence alignment was performed using Clustal Omega (Madeira *et al.,* 2024).

### Statistical analysis

Statistical analysis of all PR, LR, nodulation, IT and ethylene measurements was performed in RStudio® and are indicated in respective figure legends. Homogeneity and homoscedasticity were tested by Shapiro Wilk and Levene tests (p > 0.05) before ANOVA was performed, respectively. A Tukey post-hoc test was performed. If initial assumptions were not met, transformations (log, sqrt, Yeo-Johnson) were conducted. If assumptions were still not met, a non-parametric Scheirer-Ray-Hare or Wilcoxon test was performed and corrected for multiple comparisons with the Benjamini-Hochberg (BH) procedure. Raw data and statistical analysis results for all depicted experiments except bulk RNA-seq can be found in Supporting Information Table S1.

## Results

### LjTIE1 regulates Lotus root development and root nodule symbiosis

TIE proteins regulate primary and lateral root growth by suppressing RRB-dependent CK signaling in Arabidopsis (He *et al.,* 2022). As CK is a key regulator of both root system architecture and root nodule symbiosis, we searched for TIE homologs in Lotus. By blasting the protein sequences of the Arabidopsis homologs AtTIE1 - AtTIE4, we identified the only candidate gene *LotjaGi2glvO22O7OO* encoding a protein of 202 amino acids **(Supporting Information Figure SI).** The protein sequence contained the TIE protein-specific NLS, SPL- and EAR-motif (Tao *et al.,* 2013). Since the amino acid sequence was most similar to AtTIE1 and AtTIE2 we named the gene Lj*TIE1.* We then checked Gj*TIE1* expression in bulk RNA-seq datasets available on Lotus base (Mun *et al.,* 2016) and in our scRNA-seq datasets (Frank *et al.,* 2023) and found that Lj*TIE1* appeared to be expressed in most tissues in the root and to be regulated by CK **(Supporting Information Figure S2).** Hallmarks of an increased CK status in roots are a shortened primary root with an impaired lateral root formation (Werner & Schmülling, 2009; Del Bianco *et al.,* 2013). In legumes, external application of CK and a gain-of-function of CK signaling results in spontaneous nodulation in the absence of rhizobia and a reduced infection thread formation and nodulation capacity after rhizobia perception (Tirichine *et al.,* 2007; Heckmann *et al.,* 2011; Liu *et al.,* 2018). To validate the function of LjTIE1 in root development and nodulation, we isolated two independent *tie1* LORE presumed loss-of-function alleles **(Supporting Information Figure S3)** and evaluated their general root architecture, rhizobia-induced infection thread formation and nodulation capacity. Loss of *LjTIE1* resulted in root-level morphological changes consistent with elevated CK status. Compared to wild type, *tie1* primary roots were 60% shorter **(Figure la)** and rarely formed any lateral roots **(Figure lb, c).** Notably, both *tie1* alleles formed only 20% of nodules compared to wt **(Figure Id),** with the majority of nodules formed between 21-28 dpi rather than 7-14 dpi **(Figure Id, Supporting Information Figure S4).** Moreover, both *tie1* alleles displayed a 85 % reduction in IT density **(Figure le).** The reduced IT density phenotype paralleled the reduced nodule number, suggesting that *tie1* mutants have a specific defect in both infection and nodule organogenesis rather than only a post-infection phenotype.

**Figure 1.**
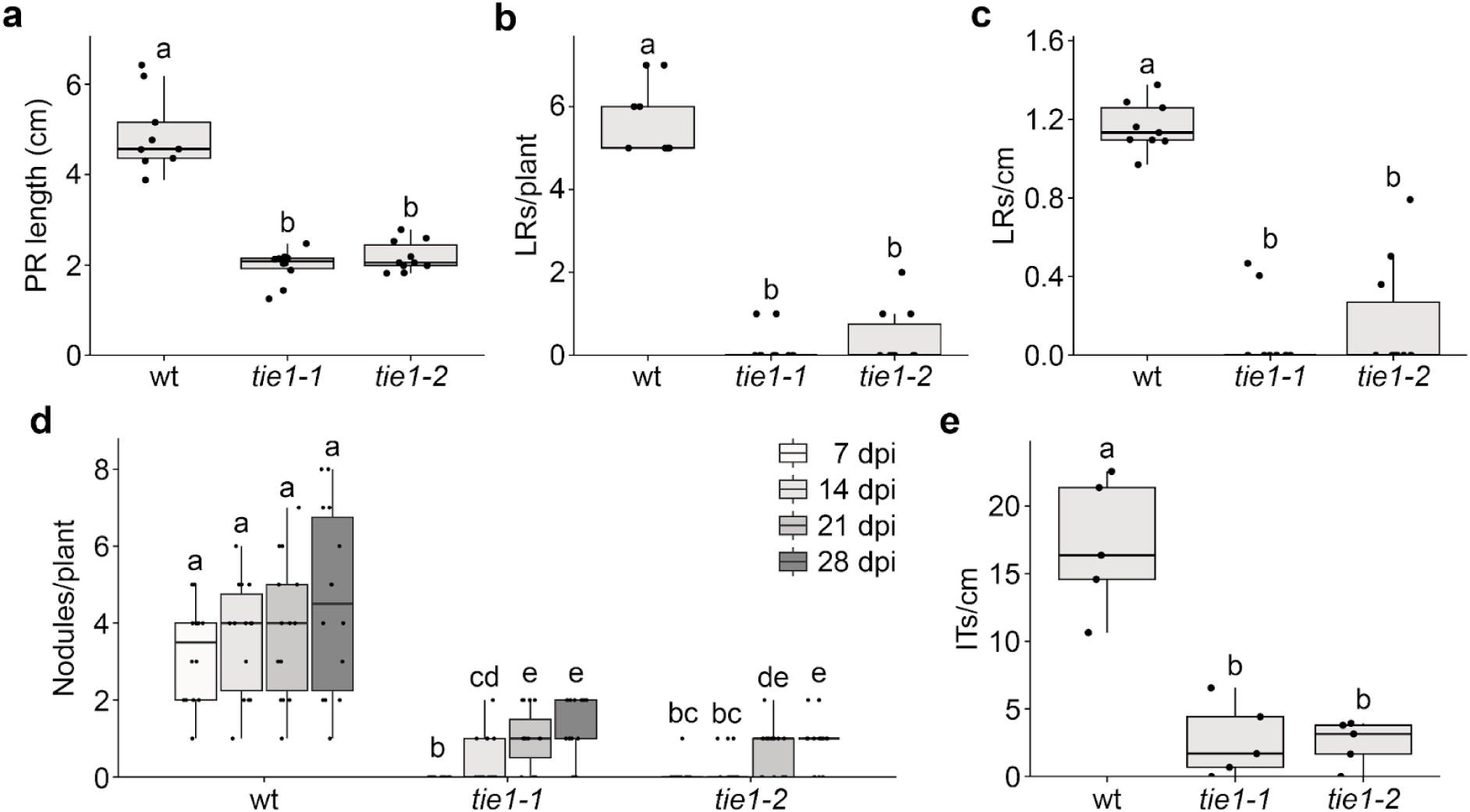
LjTIE1 is crucial for nodule organogenesis and infection. a) Primary root (PR) length, b) lateral root (LR) number and c) LR density of wt, *tie1-1* and *tie 1-2* 18 days after germination (p < 0.05; PR: one-way ANOVA followed by Tukey post-hoc test, LR/LRD: Kruskal-Wallis followed by Benjamini-Hochberg-corrected pairwise Wilcoxon rank-sum test; n > 9). d) Number of nodules at wt, *tie1-1* and *tie1-2* roots 7, 14, 21 and 28 days post inoculation (dpi; p < 0.05; Scheirer-Ray-Hare followed by Benjamini-Hochberg-corrected pairwise Wilcoxon rank-sum test; n > 14). e) Infection thread (IT) density at wt, *tie1-1* and *tie1-2* PRs 10 dpi (p < 0.05; one-way ANOVA followed by Tukey post-hoc test; n = 5). Letters indicate significantly different statistical groups.

### LjTIE1 regulates root nodule symbiosis by modulating the transcription of CK- and ethylene-related genes

Since *tie1* plants had a similar root architecture to *snf2* plants including a decreased nodule number and infection thread density after rhizobia treatment (Tirichine *et al., 2007* Liu *et al.,* 2018), we decided to study their transcriptome in comparison to each other and to wt roots. We performed a RNA-sequencing experiment of wt, *snf2, tie1-1* and *tie1-2* roots 3 dpi with either water or R7A. First, we focused on the control condition to study the base changes in the transcriptome caused by the loss of *TIE1.* Over 3300 genes were differentially expressed in at least one genotype comparison with wt and *tie1-2* having the most (1800) DEGs **(Figure 2a, Supporting Information Table S2).**

**Figure 2.**
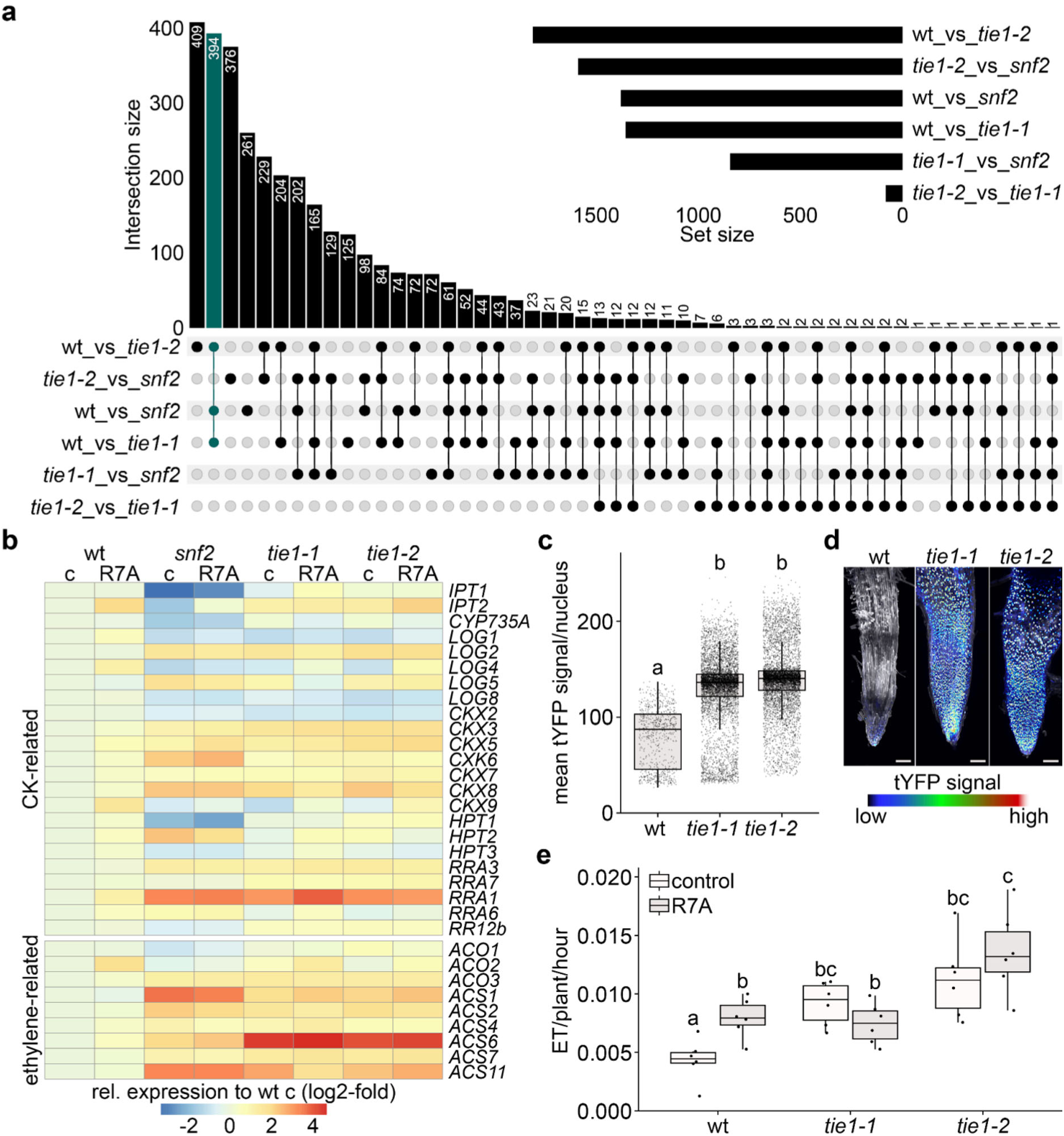
LjTIE1 modulates CK metabolism and signaling, and ethylene synthesis. a) Overlap of DEGs between wt, *snf2, tie1-1* and *tie1-2* control (c) roots. Common DEGs between the mutants compared to wt are depicted in turquoise, b) Gene expression of selected CK- and ethylene-related genes in wt, *snf2, tie1-1* and *tie1-2* roots in presence and absence of R7A 3 dpi relative to wt control, c) Mean *TCSn::tYFPnls* signal (arbitrary units) in nuclei of wt, *tie1-1* and *tie1-2* hairy roots 9 days after transfer to pots, (p < 0.05; Kruskal-Wallis followed by Benjamini-Hochberg-corrected pairwise Wilcoxon rank-sum test; n= 779 - 6093 from 5-7 hairy roots). Letters indicate significantly different statistical groups, d) Maximum projections of representative *TCSn::tYFPnls* expressing wt, *tie1-1* and *tie1-2* hairy roots 9 days after transfer to pots. All imaged hairy roots are depicted in Supporting Information Figures S5 - S7. Scale bar: 100 µm. e) Ethylene emission over 24 h for 5-day-old wt, *tie1-1* and *tie1-2* seedlings (p < 0.05; Scheirer-Ray-Hare followed by Benjamini-Hochberg-corrected pairwise Wilcoxon rank-sum test; n = 6). Letters indicate significantly different statistical groups.

As expected, the two *tie1* alleles were most similar with less than 100 DEGs. When we looked at the DEG overlaps, we found that 394 DEGs were shared between the mutants compared to the wt, which was the highest number of DEGs right after the number of DEGs between wt and *tie1-2.* Part of those 394 DEGs were genes related to CK signaling and metabolism such as *CKX, LOG* and *RRA* genes as well as *ACS* ethylene biosynthesis genes **(Figure 2b, Supporting Information Table S2).** We therefore had a closer look at CK- and ethylene-related genes being differentially expressed in at least one comparison and found that most of them showed similar trends in terms of up- or downregulation in the mutants compared to wt **(Figure 2b).** Particularly the type A response regulator *RRA1,* and multiple ethylene synthesis genes *(ACS1, ACS8, ACS IT)* were upregulated. *RRA1* induction suggests a compensatory negative feedback response to enhanced CK signaling (Imamura *et al.,* 1999). To confirm this, we checked CK signaling in wt and *tie1* hairy roots using the *TCSn::tYFPnls* reporter (Zürcher *et al.,* 2013; Reid *et al.,* 2017). Compared to wt, where tYFP signal was mostly detected in columella cells at the root tip, tYFP signal was more than 1.6-fold enhanced in *tie1* mutants and was detected throughout the whole root, indicating enhanced and broader CK signaling **(Figure 2c, d, Supporting Information Figures S5 - S7).** Next, we measured the ethylene emission of wt and *tie1* seedlings over 24 h after inoculation with water or R7A. *tie1* seedlings emitted approximately two times more ethylene than untreated wt controls **(Figure 2e).** Notably, this constitutive *tie1* ethylene emission was comparable to or exceeded the ethylene response of wt seedlings inoculated with R7A, indicating that loss of *LjTIE1* phenocopies a Nod-factor-induced ethylene burst (Reid *et al.,* 2018).

After assessing the transcriptional differences between the genotypes under control conditions, we had a closer look at the response to rhizobia. In total, 177 genes were differentially regulated in at least one genotype between control and R7A treatment. 137 DEGs were identified in wt roots upon R7A treatment of which 80 were exclusive for the wt **(Figure 3a).**

**Figure 3.**
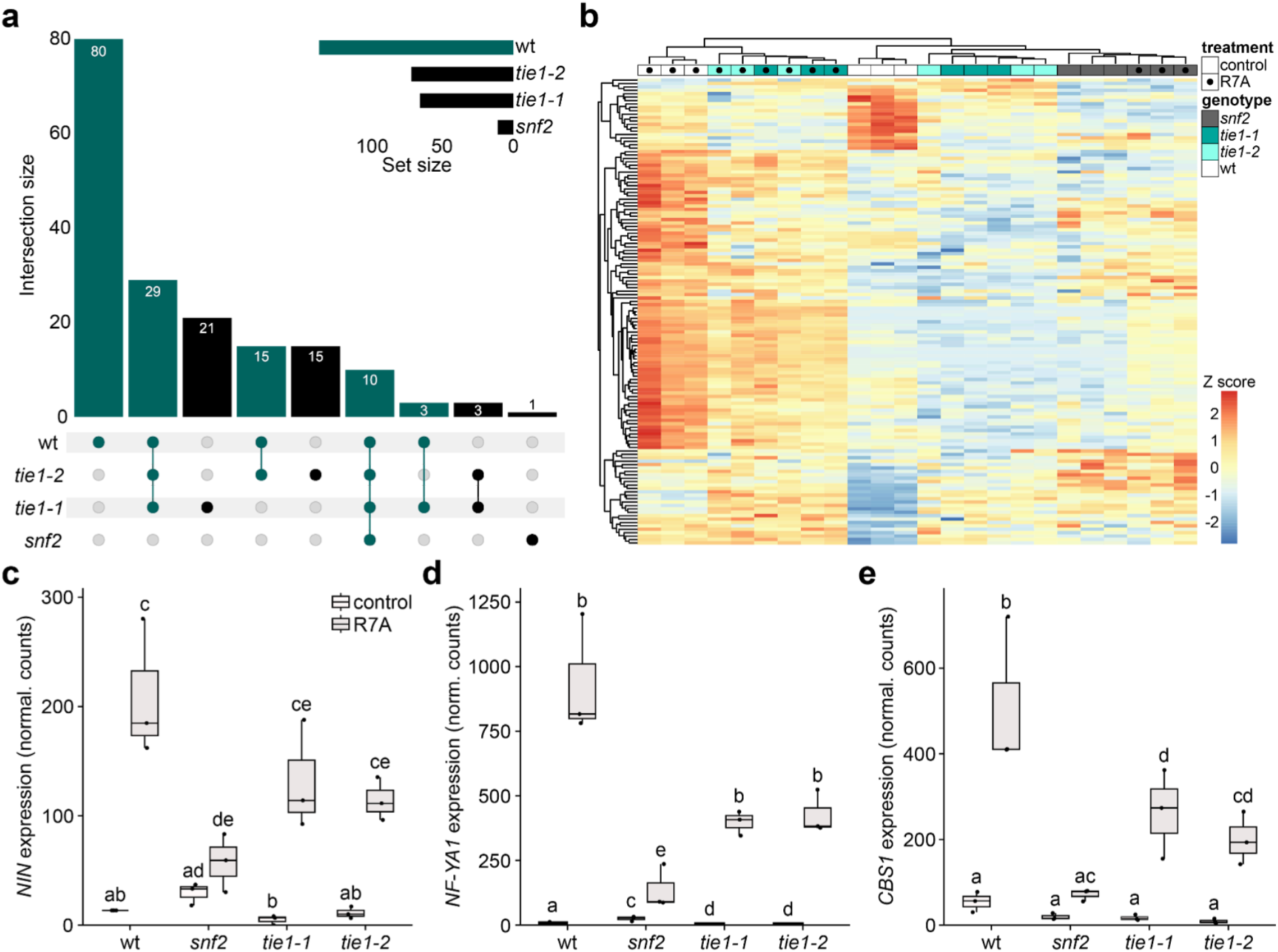
TIE1 modulates symbiotic signaling. a) Overlap of rhizobia-responsive genes between wt, *snf2, tie1-1* and *tie1-2* 3 dpi. DEGs of the wt are depicted in turquoise, b) Heatmap of genes differentially regulated upon rhizobia treatment in the wt. Gene expression is depicted as z-score in wt, *snf2, tie1-1* and *tie1-2* roots in presence and absence of R7A 3 dpi relative to wt control. Expression of a) *NIN,* b) *NF-YA1* and c) *CBS* in wt, *snf2, tie1-1* and *tie1-2* roots in the absence and presence of R7A 3 dpi (p < 0.05; two-way ANOVA followed by Tukey post-hoc test; n = 3).

Compared to that, *snf2* had only 11 exclusive DEGs, and *tie1* mutants had around 70. Strikingly, 80 out of 177 DEGs were exclusive for the wt, reflecting the differences in the inoculation and infection phenotypes between the wt and the mutants. We then used all 137 wt-related DEGs indicating functional nodule organogenesis and IT formation for our clustering analysis **(Figure 3b).** The analysis revealed that wt, *tie1-1* and *tie1-2* R7A samples formed their own cluster suggesting a similar response to rhizobia. *snf2* control and R7A samples formed their own subcluster within the remaining samples reflecting the impairment in rhizobia-induced symbiosis signaling. Some of the induced nodulation-related genes such as *NF-YA1* and *NIN* were elevated in *snf2* compared to the other genotypes under control conditions, indicating a rhizobia-independent activation of the LHK1-dependent symbiosis pathway **(Figure 3c, d).** Moreover, while *tie1* plants responded stronger to rhizobia than *snf2,* their response was dampened compared to wt reflecting the lower nodule and IT number in those mutants **(Figure 3b - e).**

### Inhibition of ethylene synthesis rescues *tie1* nodulation

As *tie1* plants emitted more ethylene we were wondering whether the observed phenotypes could be rescued by preventing ethylene biosynthesis. Like *tie1* plants, the CK gain-of-function mutant *snf2* is impaired in nodulation and IT formation, and displays increased ethylene emission (Reid *et al.,* 2018). Inhibiting ethylene biosynthesis in *snf2* through AVG rescues those phenotypes and enhances the spontaneous nodulation phenotype in the absence of rhizobia (Liu *et al.,* 2018). We therefore cultivated *tie1* plants on AVG plates and quantified nodulation and infection threads. On AVG plates, *tie1* roots formed significantly more nodules compared to control conditions 14 dpi and AVG-supplemented *tie1* roots formed 20 % more nodules as AVG-supplemented wt roots **(Figure 4a, b).**

**Figure 4.**
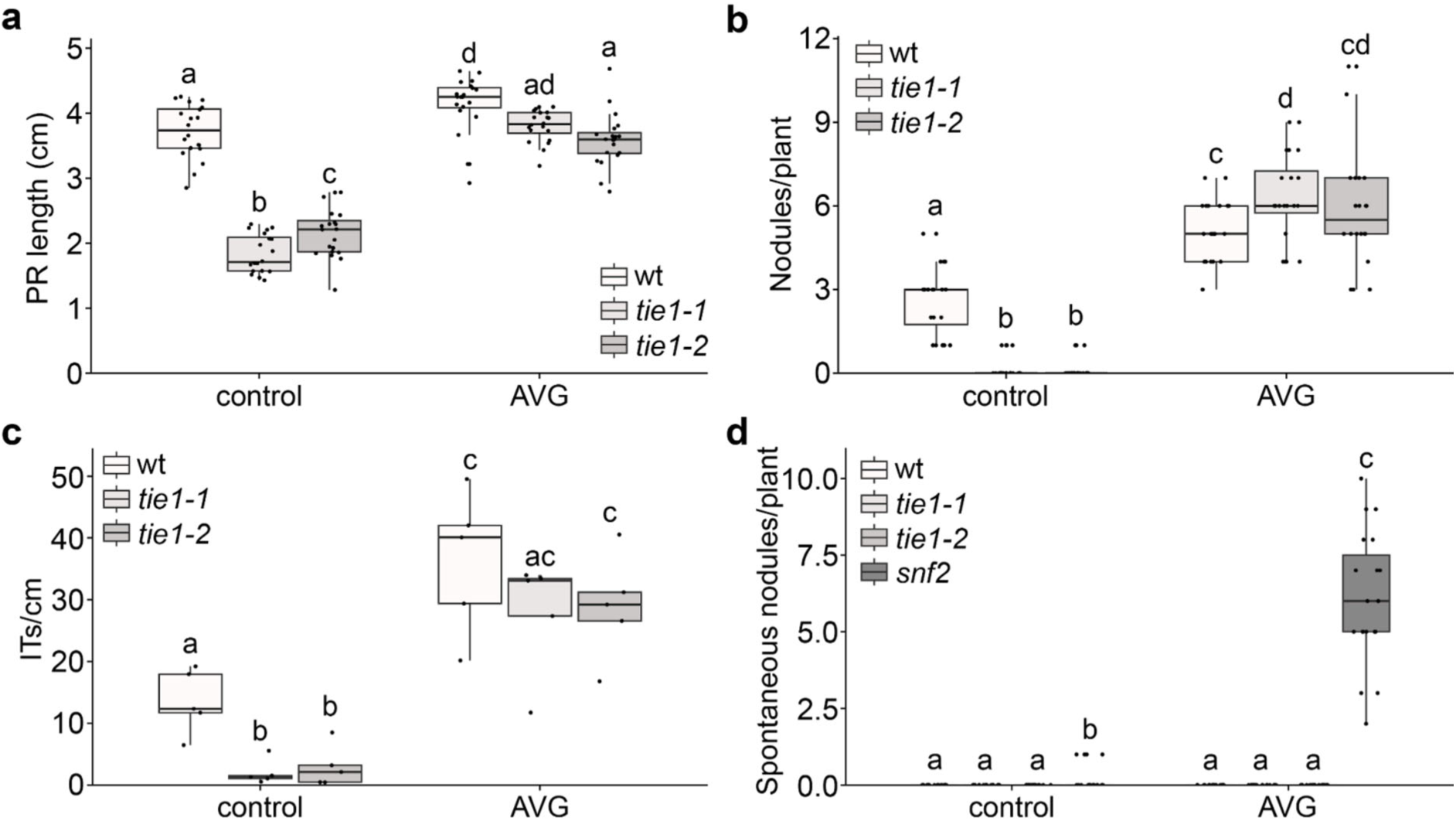
Impairment of ethylene synthesis rescues the loss of *LjTIE1.* a) Primary root (PR) length of and b) number of nodules at wt, *tie1-1* and *tie1-2* roots in the absence and presence of 1 µM aminoethoxyvinylglycine (AVG) 14 dpi (p < 0.05; PR: two-way ANOVA followed by Tukey post-hoc test, nodules: Scheirer-Ray-Hare followed by Benjamini-Hochberg-corrected pairwise Wilcoxon rank-sum test; n = 20). c) Infection thread (IT) density at wt, *tie1-1* and *tie1-2* roots 10 dpi in the absence and presence of 1 µM AVG (p < 0.05; two-way ANOVA followed by Tukey post-hoc test; n = 5). d) Spontaneous nodules at wt, *tie1-1, tie1-2* and *snf2* roots five weeks after germination in the absence and presence of 1 µM AVG (p < 0.05; Scheirer-Ray-Hare followed by Benjamini-Hochberg-corrected pairwise Wilcoxon rank-sum test; n > 19). Letters indicate significantly different statistical groups.

Most of the nodules at the wt under control conditions were pink, while AVG treatment reduced the number of pink nodules and induced the formation of white nodules **(Supporting Information Figure S8).** Compared to wt, *tie1* roots formed more pink nodules when AVG was applied and the number of pink nodules was similar to that of wt under control conditions **(Supporting Information Figure S8b).** We then quantified ITs and found that the reduced IT density at *tie1* roots was rescued by AVG **(Figure 4c).** Lastly, we assessed the spontaneous nodulation capacity of *tie1* mutants to that of *snf2,* as *tie1* phenocopied *snf2* in all other parameters that are indicative of an enhanced CK status. To our surprise, wt and *tie1* roots did not form any spontaneous nodules in the absence and presence of AVG five weeks after germination, while *snf2* formed spontaneous nodules under both conditions **(Figure 4d).** We hypothesized that a nodulation-specifíc CK stimulus - beyond the constitutively enhanced CK perception in *tie1* roots - was required to trigger nodule organogenesis. To test this, we applied the synthetic CK BAP in combination with AVG to wt and *tie1* seedlings. This dual treatment induced robust spontaneous nodulation and revealed a *tie1* hypernodulation phenotype **(Figure 5a).** At 14 dpi, *tie1-1* and *tie1-2* formed 2.1- and 2.3-fold more nodules than wt, respectively. However, by 21 dpi, nodule numbers in *tie1-1* roots had converged to wt levels, whereas *tie1-2* maintained a 1.4-fold higher nodule number relative to wt through 28 dpi **(Figure 5a),** suggesting possible mild allele-specific effects. In addition to the spontaneous nodules phenotype, both alleles formed additional epidermal/cortical tissue indicating additional nodulation-unrelated CK-hypersensitivity **(Figure 5b).**

**Figure 5.**
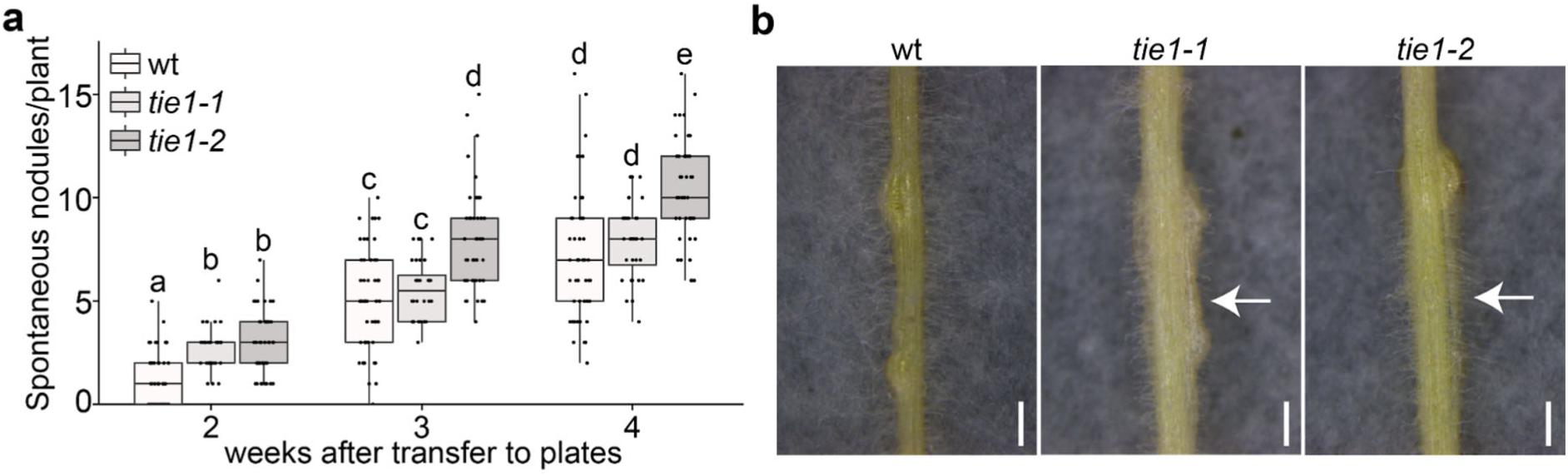
A loss of TIE1 causes hyper-sensitivity to BAP. a) Quantification of spontaneous nodules at wt, *tie1-1* and *tie1-2* roots two to four weeks after transfer to plates containing 1 µM AVG and 10 nM BAP (p < 0.05; Scheirer-Ray-Hare followed by Benjamini-Hochberg-corrected pairwise Wilcoxon rank-sum test; n > 32). Letters indicate significantly different statistical groups, b) Pictures of representative roots with spontaneous nodules and excessive tissue (white arrows) 28 dpi. Scale bar: 1mm.

## Discussion

Our characterization of LjTIE1 revealed that legumes employ multiple mechanisms to constrain CK signaling during root nodule symbiosis, preventing constitutive nodulation and coordinating the spatial and temporal progression of infection and organogenesis. Three key findings support this conclusion: First, *tie1* loss-of-function mutants exhibit constitutively elevated CK signaling yet fail to form spontaneous nodules, demonstrating that CK signal intensity alone is insufficient for autonomous organogenesis. Second, *tie1* phenotypes - including impaired root growth, reduced lateral root density, and decreased nodulation - are fully rescued by ethylene biosynthesis inhibition, revealing that LjTIE1 functions primarily by preventing CK-induced ethylene overproduction. Third, despite lacking spontaneous nodulation, *tie1* mutants hypernodulate in response to exogenous CK application, indicating that they retain competence for CK-driven organogenesis but require supraoptimal CK levels to overcome altered regulatory thresholds. Together, these results establish LjTIE1 as a critical rheostat that dampens CK signaling after symbiotic induction, enabling precise spatiotemporal control of nodule development.

### LjTIE1 functions as a CK-inducible negative regulator

The molecular mechanism by which LjTIE1 attenuates CK signaling likely parallels the characterized Arabidopsis TIE function. In Arabidopsis roots, AtTIE1 and AtTIE2 are transcriptionally induced by CK and establish negative feedback by recruiting TPL/TPR co-repressors to RRBs, thereby inhibiting CK-responsive gene expression (He *et al.,* 2022). Our data support conserved function in Lotus: *LjTIE1* is upregulated by both CK and Nod factor **(Supporting Information Figure S2),** *tie1* mutants show elevated *TCSn* reporter activity throughout roots tissues and differential expression of CK-responsive genes **(Figure 2),** confirming that TIE1 normally dampens CK signaling output.

Notably, both *tie1* and *snf2* mutants show strong upregulation of *RRA1* **(Figure 2b),** a RRA that functions as a primary negative feedback component in CK signaling (Brandstatter & Kieber, 1998). RRAs are rapidly induced by CK and attenuate signaling by competing with RRBs for phosphotransfer from HPTs (To *et al.,* 2004). The parallel *RRA1* induction in *tie1* and *snf2 -* despite their mechanistically distinct origins of elevated CK signaling (loss of downstream negative regulation versus constitutive receptor activation, respectively) - indicates that both mutants trigger the canonical CK negative feedback pathway. However, *RRA1* upregulation alone is evidently insufficient to fully compensate for *LfHEl* loss or *snf2* receptor hyperactivation, suggesting that LjTIE1 and RRAs operate through complementary mechanisms to constrain CK responses. LjTIE1 likely provides RRB-level transcriptional repression, while RRAs function at the phosphorelay level - together creating layered negative feedback that fine-tunes CK signaling intensity and duration.

### Why *tie1* mutants lack spontaneous nodulation

The absence of spontaneous nodulation in *tie1* **(Figure 4)** is our most surprising finding and gives critical insights into the onset of nodulation. Gain-of-function mutations in the CK receptor LHK1 - such as *snf2 (Tirichine *et al.,* 2007)* or *snf5 (Liu *et al.,* 2018)* - consistently trigger nodule formation without rhizobia by causing constitutive LHK1 signaling. Given that *tie1* exhibited elevated CK signaling comparable to *snf2* (evidenced by enhanced TCSn reporter intensity and similar regulation of CK-responsive genes including *RRA1* **[Figure 2]),** the absence of spontaneous nodulation demands explanation.

We propose that the critical distinction lies in the receptor source and signaling quality of elevated CK responses. The *snf2* mutation causes constitutive activation specifically of LHK1 (Tirichine *et al.,* 2007), providing sustained LHKl→RRB→NIN signaling that mimics authentic symbiotic induction. In contrast, LjTIE1 loss removes negative regulation downstream of receptor activation, likely affecting signaling through all four Lotus histidine kinases (LHK1, LHK1 A, LHK2, LHK3) that show partially redundant functions (Held *et al.,* 2014). While LHK1 is essential for nodulation, LHK1A, LHK2 and LHK3 contribute primarily to lateral root development and general CK responses. Thus, *tie1’s* pan-receptor CK elevation may enhance general root CK responses without achieving the LHK1-specific signal intensity required to cross the threshold for nodule organogenesis.

Supporting this receptor-specificity model, *tie1* and *snf2* transcriptomes differed substantially beyond their shared upregulation of canonical CK-responsive genes, suggesting qualitatively different signaling outputs despite both showing elevated CK signaling. Additionally, *tie1* mutants showed no altered *LHK1* expression in our bulk RNA-seq data, indicating elevated signaling arises from impaired negative regulation rather than altered receptor abundance. The tissue distribution of TCSn signal in *tie1* appeared broadly elevated rather than cortex-specific **(Figure 2c, d),** whereas spontaneous nodulation requires cortex-enriched CK signaling (Tirichine *et al.,* 2007; Reid *et al.,* 2017).

Alternative explanations include insufficient CK signaling intensity despite elevation (testable by e.g. crossing *tie1* and *snf2* mutants), inappropriate developmental timing of CK elevation, or requirements for rhizobial signals beyond CK (Yang *et al.,* 2022). The observation that AVG treatment - which eliminates elevated ethylene - does not induce spontaneous nodules in *tie1* **(Figure 4d)** argues against ethylene being the sole barrier. Single-cell transcriptomics comparing epidermal versus cortical gene expression in *tie1* would test whether elevated CK signaling occurs uniformly or shows tissue-specific patterns invisible to whole-root analyses.

### LjTIE1’s role in fine-tuning CK-ethylene crosstalk

The complete rescue of *tie1* phenotypes by ethylene biosynthesis inhibition **(Figure 4)** reveals that LjTIE1’s regulatory function operates primarily through modulating CK-ethylene crosstalk. This crosstalk is well-established: elevated CK induces ethylene biosynthesis genes *(ACS, ACO),* and resulting ethylene accumulation inhibits root elongation and lateral root formation (Ruzicka *et al.,* 2009; He *et al.,* 2022; Yamoune *et al.,* 2024). Our transcriptomic data revealed significant upregulation of ethylene biosynthesis genes in *tie1* **(Figure 2b),** and quantitative measurements confirmed a 2-fold elevated emission **(Figure 2e),** suggesting that LjTIE1 inhibits ethylene production.

The molecular mechanism likely involves transcriptional regulation: RRBs directly activate *ACS* and *ACO* gene expression in response to CK (Yamoune *et al.,* 2024), and by attenuating RRB activity through corepressor recruitment, LjTIE1 dampens this CK-to-ethylene transcriptional cascade. The functional significance extends specifically to nodule organogenesis. Ethylene plays stage-specific roles in legume-rhizobia symbiosis: early infection requires an ethylene spike, but sustained high ethylene inhibits both infection thread progression and nodule organogenesis (Reid *et al.,* 2018; Lin *et al.,* 2020). Enhanced ethylene in *tie1* mutants likely explains reduced infection threads and nodule numbers **(Figure 1),** as elevated ethylene chronically suppresses both processes. The rescue of its nodulation and infection phenotype under exogenous AVG **(Figure 4)** phenocopying *snf2* (Reid *et al.,* 2018; Liu *et al.,* 2018) supports this interpretation.

LjTIE1’s role in CK-ethylene crosstalk may also connect to systemic autoregulation of nodulation (AON). While ethylene inhibits nodulation locally in the first cortex layer opposite the phloem pole (Heidstra *et al.,* 1997), CK integrates local and systemic nodulation signals (Sasaki *et al.,* 2014). If LjTIE1 is required to interpret CK signaling intensity as a proxy for nitrogen status (Cortleven *et al.,* 2019; Lin *et al.,* 2021), it might suppress nodulation systemically under nitrogen-sufficient conditions by directly altering CK signaling or indirectly through ethylene accumulation. Testing *tie1* nodulation responses and *LjTlE1* expression across nitrogen gradients would clarify whether LjTIE1 acts as an integrator of nutritional and symbiotic signaling.

### Spatial and temporal control of symbiotic CK responses

A fundamental question is how CK drives opposing outcomes in adjacent tissues: promoting cortical organogenesis while suppressing epidermal infection. LjTIE1 may contribute to this spatial specificity through differential expression or activity between tissues, tissue-specific RRB interactions, or differential corepressor availability in cortical nodule primordia. Single-cell RNA-seq data supports this view as Lj*TIE1* was not expressed ubiquitously, and bulk RNA-seq showed its induction by CK and rhizobia perception within 24 h **(Supporting Information Figure S2).** Notably, our TCSn reporter data displayed broad elevation in *tie1* **(Figure 2c, d),** suggesting LjTIE1 regulates CK signaling generally rather than tissue-specifically. However, steady-state imaging may miss transient, tissue-specific functions. Single-cell RNA-seq comparing epidermal with cortical transcriptomes in wt versus *tie1* during active nodulation would test whether LjTIE1 preferentially regulates CK responses in specific tissues.

Beyond spatial control, LjTIE1 may provide temporal gating. As a CK-inducible negative regulator, LjTIE1 establishes negative feedback: initial Nod-factor-induced CK activates LHK1→RRBs→nodulation genes but simultaneously induces *LjTIE1,* which subsequently dampens RRB activity. This feedback mechanism creates transient CK signaling pulses rather than sustained activation. Recent work has demonstrated that such pulsatile CK dynamics are functionally critical—sustained high CK signaling inhibits both infection thread formation and nodule organogenesis, whereas precisely timed CK pulses promote symbiotic development (Soyano *et al.,* 2024). This temporal specificity may explain why constitutive CK signaling through receptor gain-of-function mutations causes ectopic developmental outcomes, while feedback-regulated pulsatile signaling drives appropriate organogenesis. The LjTIE1-mediated negative feedback loop would ensure that initial CK peaks triggering nodule initiation are followed by signal attenuation, preventing the sustained high CK levels that Soyano et al. (2024) showed to be inhibitory. Time-course experiments measuring Lj*TIE1* expression, CK levels, and ethylene emission at hourly intervals during the first 48 hours post-inoculation would test this temporal model.

#### Working Model

Integrating our findings, we propose that the perception of rhizobia via Nod factors activates symbiotic signaling resulting in CK synthesis **(Figure 6).** CKs are perceived by LHK1, triggering phosphotransfer to RRBs. Activated RRBs drive nodule organogenesis by inducing the transcription of downstream genes such as *NIN,* while simultaneously inducing Lj*TIE1* transcription. LjTIE1 recruits TPL/TPR corepressors to RRBs, attenuating transcriptional activity and constraining CK signaling. This feedback prevents excessive ethylene accumulation and limits spatial-temporal CK response extent. In *tie1* mutants, loss of feedback causes constitutive RRB activity, chronic ethylene overproduction inhibiting root growth and nodulation, and altered CK response thresholds. Despite elevated CK signaling, *tie1* lacks spontaneous nodulation because elevated signaling affects multiple receptors rather than providing LHK1-specific activation that *snf2* achieves (Tirichine *et al.,* 2007).

**Figure 6.**
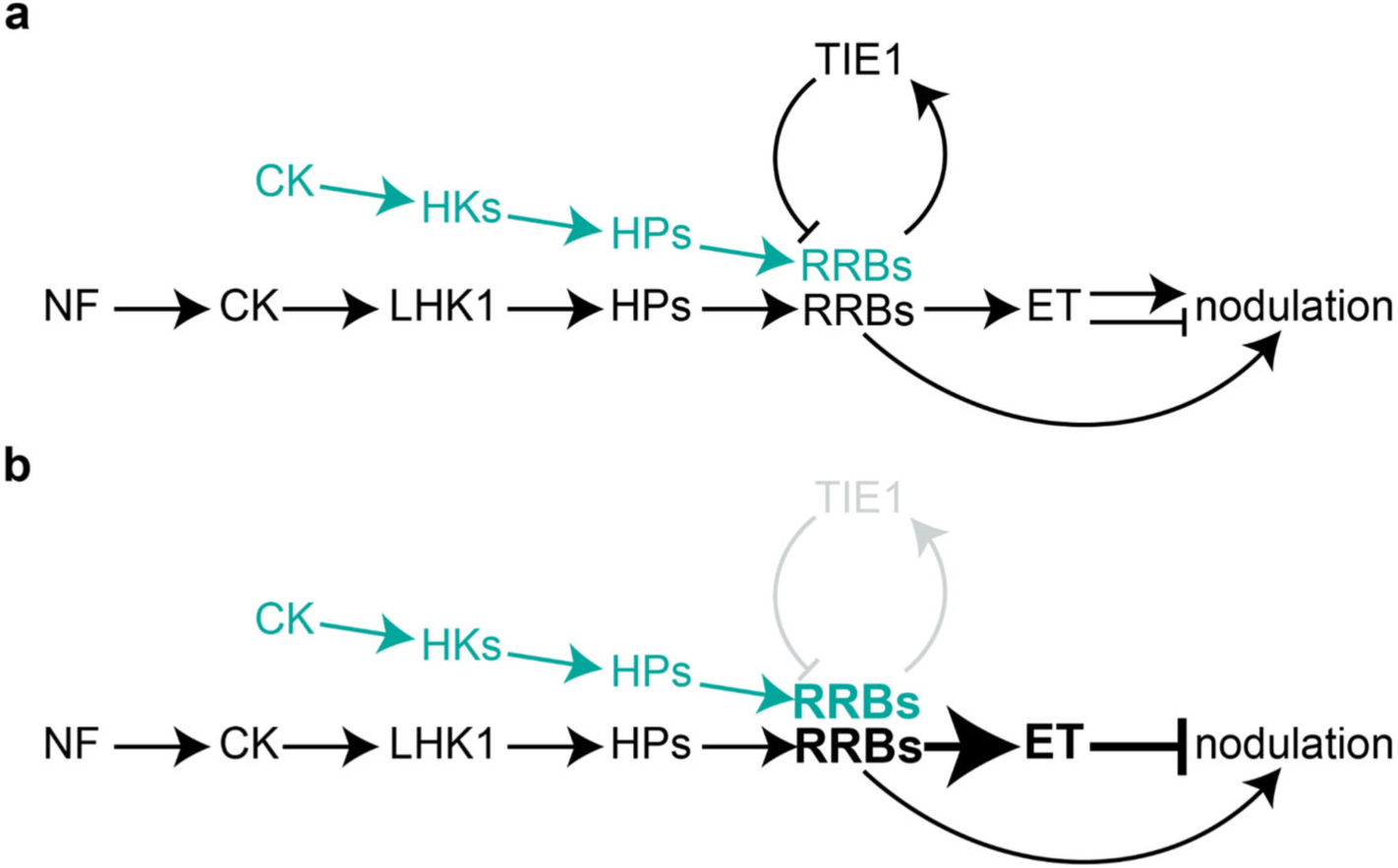
Model of CK- and LjTIE1-dependent root nodule symbiosis in Lotus. a) Upon rhizobia-derived Nod factor (NF) perception, CK is induced and triggers nodule organogenesis mainly through CK receptor LHK1 and ethylene (ET) synthesis. LjTIE1 balances LHK1-dependent and LHK1-independent signaling (turquoise) by negatively regulating RRB activity. This balanced CK signaling enables nodulation. b) In the absence of LjTIE1, RRBs are not inhibited resulting in chronic ET overproduction inhibiting nodulation.

### Limitations and Broader Implications

While TCSn reporter analysis demonstrates elevated CK signaling **(Figure 2),** we have not directly measured CK levels - enhanced signals could reflect increased abundance, receptor sensitivity, or prolonged RRB activity. LC-MS/MS quantification would distinguish these. Our ethylene measurements **(Figure 2)** do not resolve tissue-specific production, which may differentially affect infection versus organogenesis. The divergent kinetics between *tie1-1* and *tie1-2* at 21 dpi **(Figure 5a)** requires further investigation for true allele-specificity versus stochastic variation.

Our identification of LjTIE1 as a negative regulator illuminates how legumes achieve developmental specificity during symbiosis. Negative regulators modulating CK output provide mechanisms for tissue-specific responses where identical signals drive different outcomes. From an evolutionary perspective, recruitment of CK negative regulators to control symbiosis reflects co-option of pre-existing root developmental modules - TIE proteins regulate CK-dependent root development in non-legumes (He *et al.,* 2022), suggesting legumes adapted existing circuits when symbiosis evolved. Understanding which components are symbiosis-specific versus borrowed from root programs would illuminate genetic innovations underlying symbiosis evolution and inform engineering efforts for non-legume crops.

In conclusion, LjTIE1 provides negative feedback that balances CK signaling during symbiosis, preventing constitutive nodulation and coordinating infection with organogenesis. By linking CK perception to ethylene biosynthesis and constraining signaling intensity, LjTIE1 enables precise spatiotemporal control required for successful nitrogen-fixing symbiosis.

## Acknowledgements

This work was supported by the European Research Council (ERC) under the European Union’s Horizon 2020 research and innovation programme (grant agreement no. 834221) and by the project Enabling Nutrient Symbioses in Agriculture (ENSA), that is funded by Gates Agricultural Innovations (INV-57461). The authors thank Stig U. Andersen and Jens Stougaard for their financial support, Dugald Reid for giving feedback on the manuscript, and Antje von Schaewen for recommending the first to the corresponding author.

## Author contributions

ML and MF performed experiments, analyzed and interpreted data. MF conceptualized the research and wrote the manuscript. ML and MF edited the manuscript.

## Data availability

The data that support the findings of this study are available in the Supporting Information of this article. Sequencing raw data is deposited at the Sequence Read Archive (SRA4) with the BioProject ID PRJNA1333668.

---

**Figure S1.**
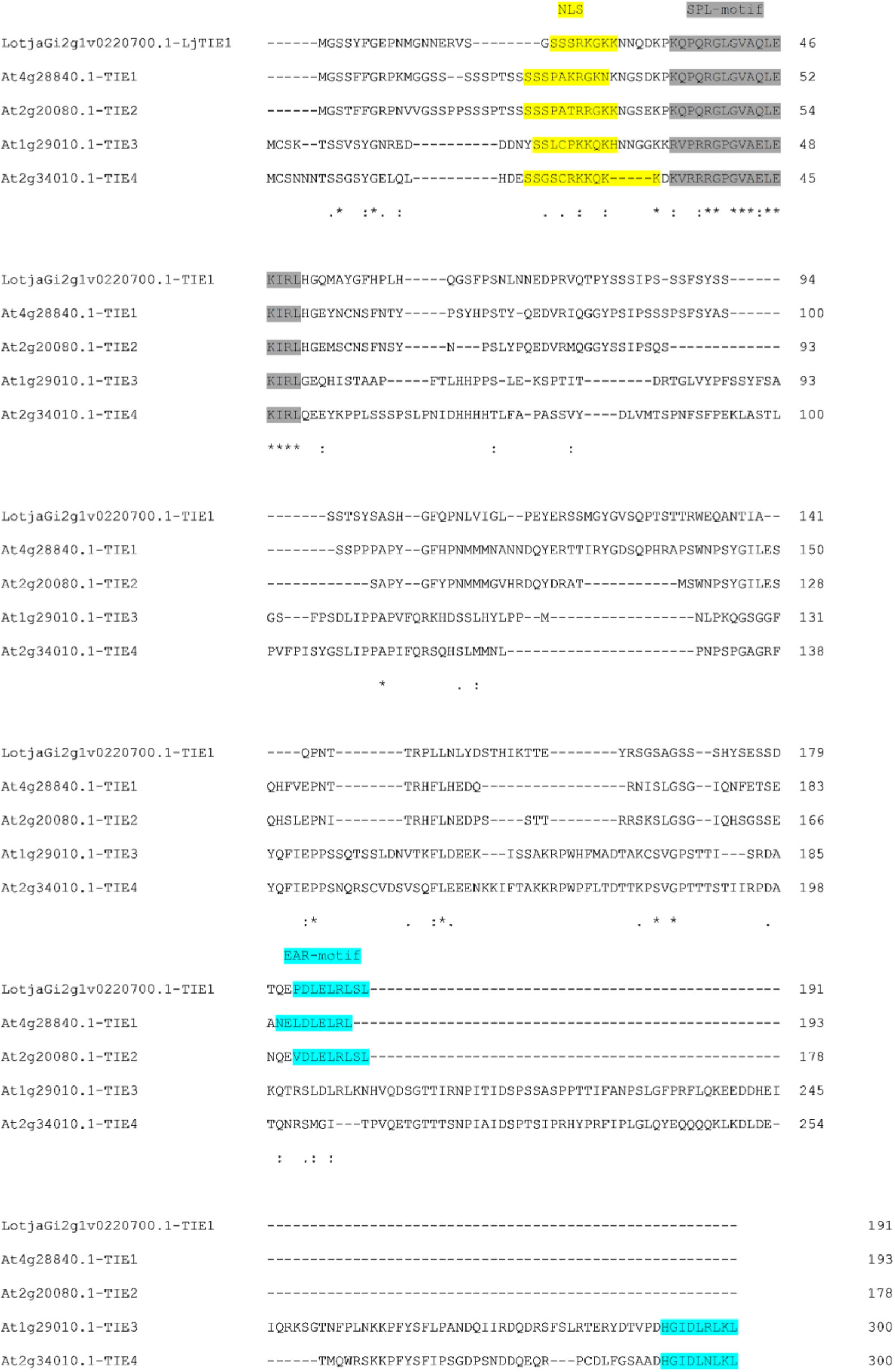
LjTIE1 shares homology to Arabidopsis AtTIE1 and AtTIE2. Amino acid sequence alignment of LjTIE1 and AtTIEs. Highlighted are the nuclear localization signal (NLS, yellow), SPL- and EAR-motives (grey and cyan).

**Figure S2.**
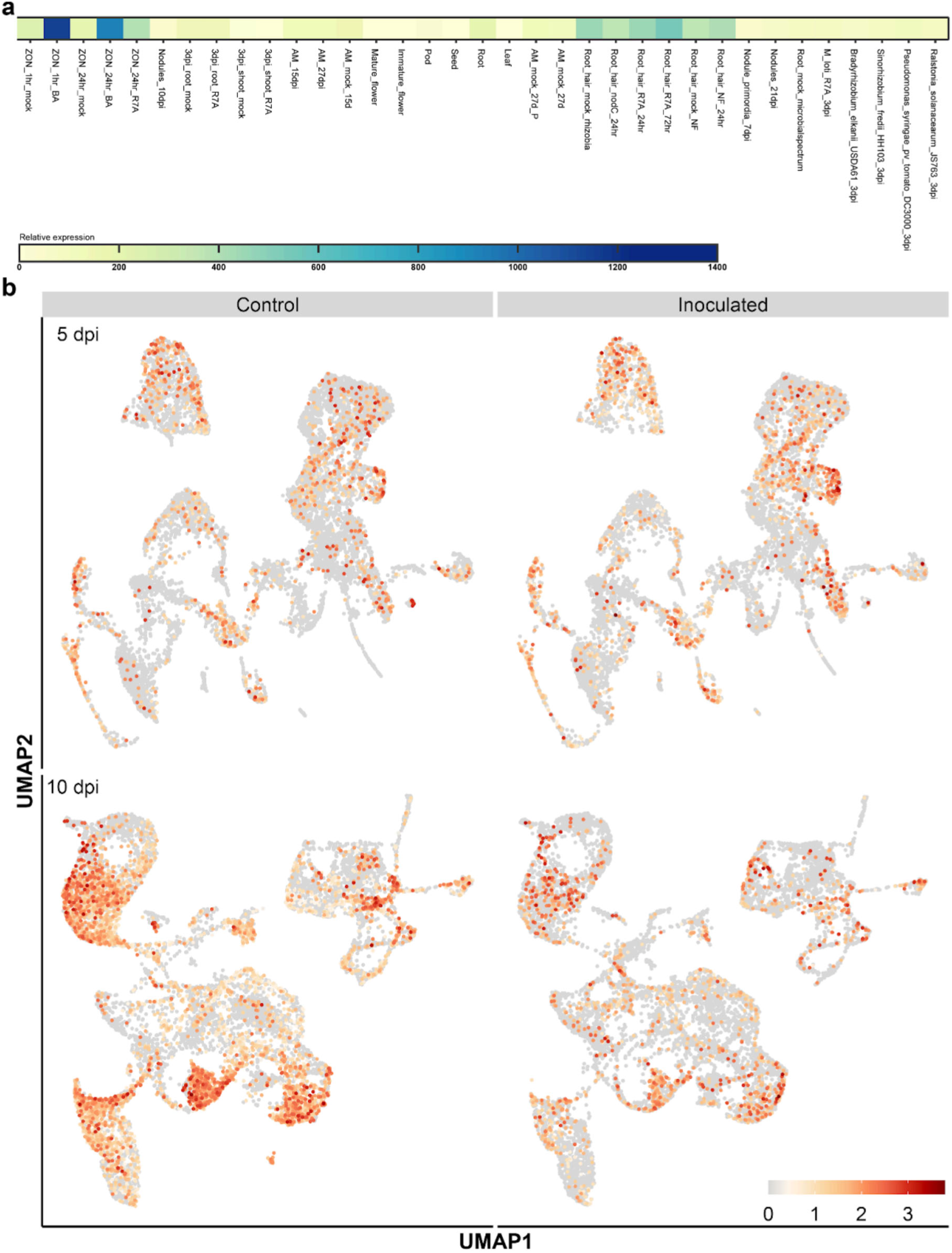
Expression of *LjTIE1* in published bulk and single-cell RNA-seq data. a) Relative expression of *TIE1* in bulk RNAseq data on Lotus base, b) Relative expression of *TIE1* in Lotus roots 5 and 10 dpi.

**Figure S3.**
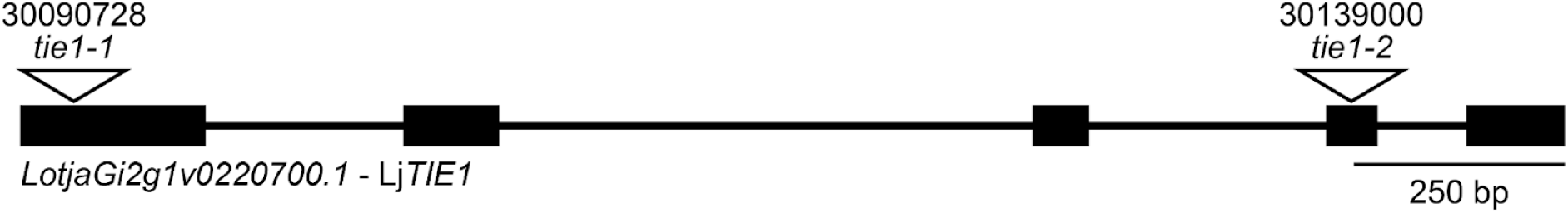
Overview over *LjTIE1* and *tie1* alleles. Rectangles represent exons and lines represent introns. Numbers above alleles represent LORE lines used (Malolepszy *et al.,* 2016).

**Figure S4.**
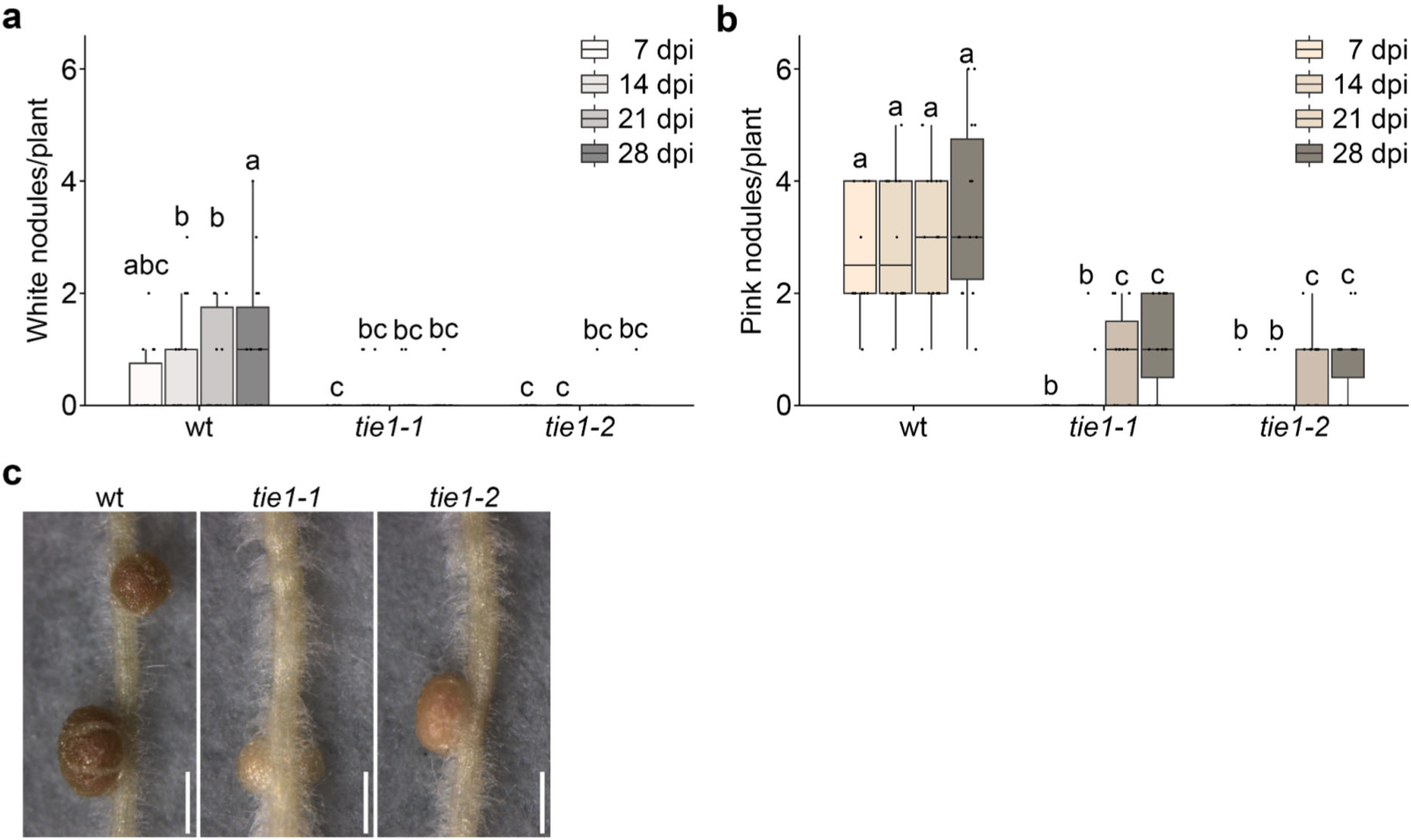
White and pink nodules at *tie1* roots. a) Number of white and b) of pink nodules at wt, *tie1-1* and *tie1-2* roots 7, 14, 21 and 28 days post inoculation (dpi; p < 0.05; Scheirer-Ray-Hare with Benjamini-Hochberg-corrected Wilcoxon rank sum test; n > 14). Letters indicate significantly different statistical groups, c) Pictures of representative roots with nodules 28 dpi. Scale bar: 1mm.

**Figure S5.**
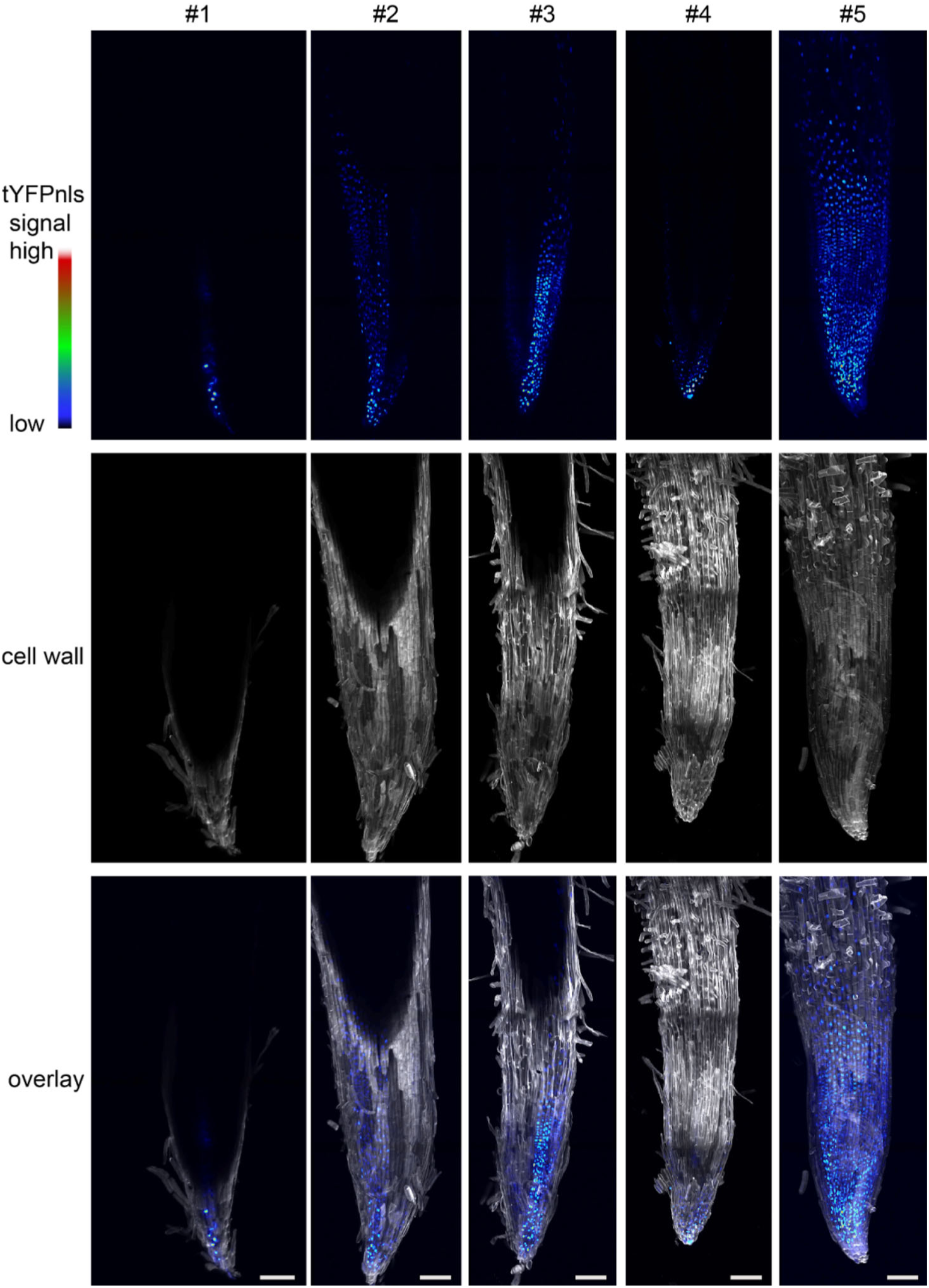
*TCSn::tYFPnls* expression in wt transgenic hairy roots. Maximum projections of all imaged wt hairy roots expressing the *TCSn::tYFPnls* construct for the experiment performed in Figure 2. Scale bar: 100 µm.

**Figure S6.**
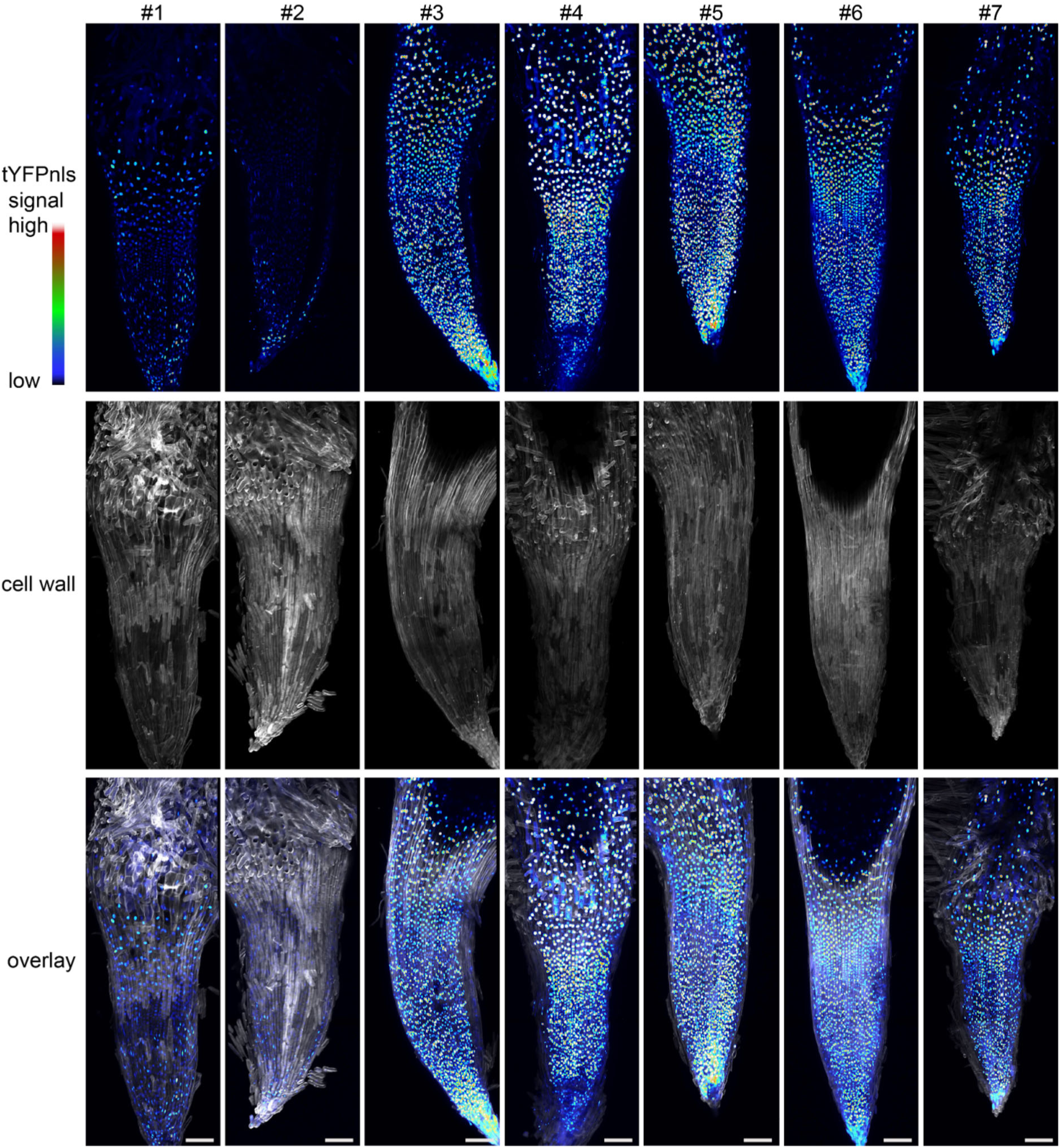
*TCSn::tYFPnls* expression in *tie1-1* transgenic hairy roots. Maximum projections of all imaged *tie1-1* hairy roots expressing the *TCSn::tYFPnls* construct for the experiment performed in Figure 2. Scale bar: 100 µm.

**Figure S7.**
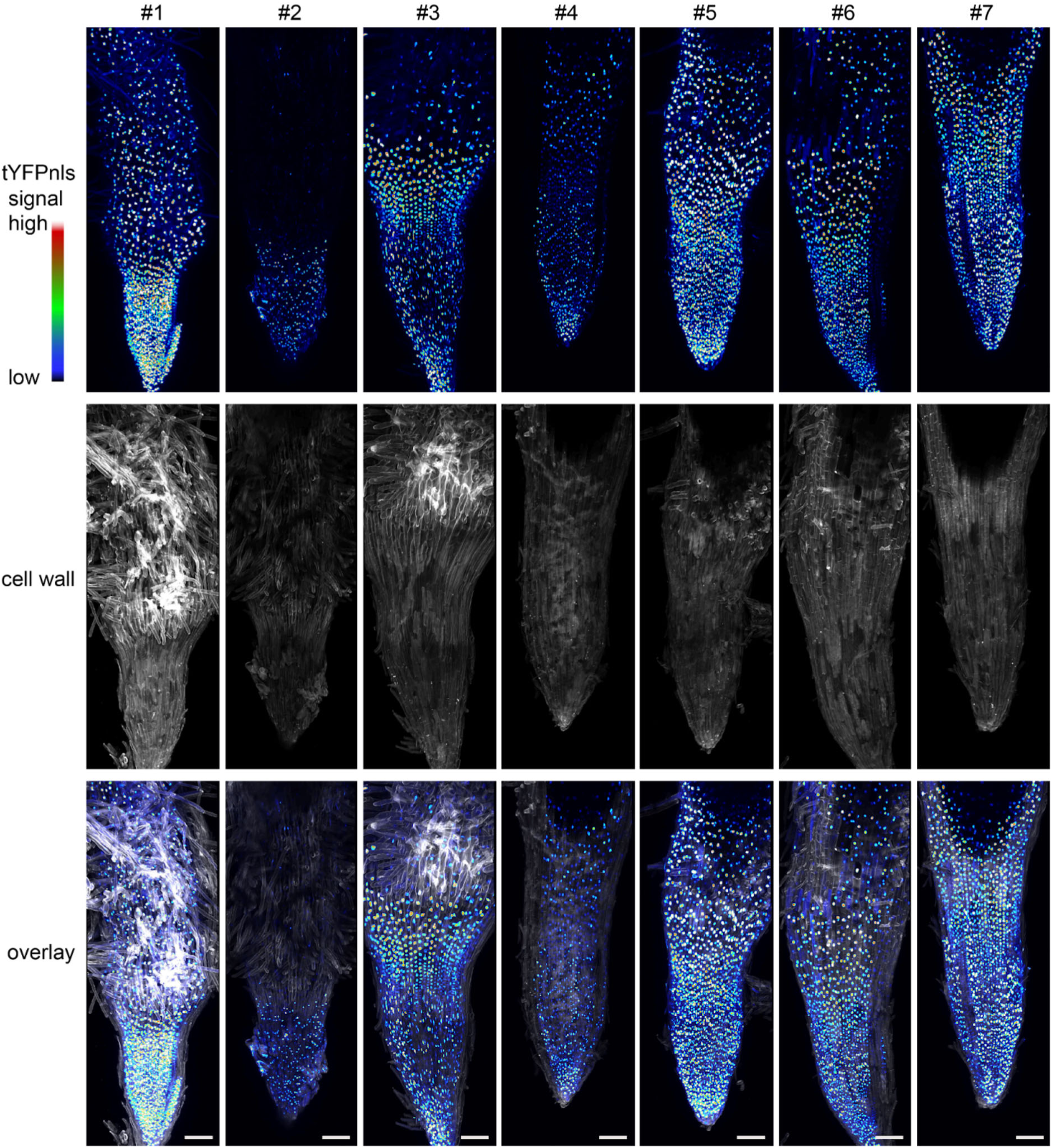
*TCSn::tYFPnls* expression in *tie1-2* transgenic hairy roots. Maximum projections of all imaged *tie1-2* hairy roots expressing the *TCSn::tYFPnls* construct for the experiment performed in Figure 2. Scale bar: 100 µm.

**Figure S8.**
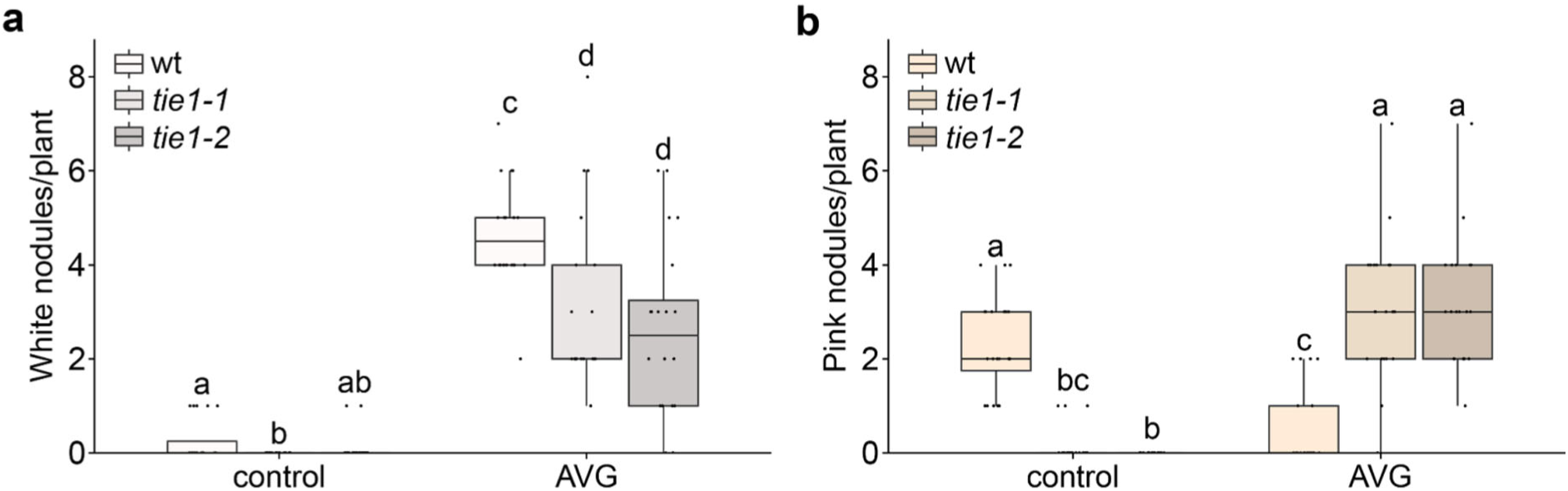
White and pink nodules at *tie1* roots in presence and absence of AVG. Number of a) white and b) pink nodules at wt, *tie1-1* and *tie1-2* roots in the absence and presence of 1 µM aminoethoxyvinylglycine (AVG) 14 dpi (p < 0.05; white: Scheirer-Ray-Hare with Benjamini-Hochberg-corrected Wilcoxon rank sum test; n = 20). Letters indicate significantly different statistical groups.

